# iEnhancer-CLA: Self-attention-based interpretable model for enhancers and their strength prediction

**DOI:** 10.1101/2021.11.23.469658

**Authors:** Lijun Cai, Xuanbai Ren, Xiangzheng Fu, Mingyu Gao, Peng Wang, Junling Xu, Wei Liu, Zejun Li, Xiangxiang Zeng

**Affiliations:** College of Computer Science and Electronic Engineering, Hunan University, Hunan Changsha, China; College of Computer Science, Xiangtan University, Xiangtan, China; College of Computer and Information Science, Hunan Institute of Technology, Hengyang, China

## Abstract

Enhancer is a class of non-coding DNA cis-acting elements that plays a crucial role in the development of eukaryotes for their transcription. Computational methods for predicting enhancers have been developed and achieve satisfactory performance. However, existing computational methods suffer from experience-based feature engineering and lack of interpretability, which not only limit the representation ability of the models to some extent, but also make it difficult to provide interpretable analysis of the model prediction findings.In this paper, we propose a novel deep-learning-based model, iEnhancer-CLA, for identifying enhancers and their strengths. Specifically, iEnhancer-CLA automatically learns sequence 1D features through multiscale convolutional neural networks (CNN), and employs a self-attention mechanism to represent global features formed by multiple elements (multibody effects). In particular, the model can provide an interpretable analysis of the enhancer motifs and key base signals by decoupling CNN modules and generating self-attention weights. To avoid the bias of setting hyperparameters manually, we construct Bayesian optimization methods to obtain model global optimization hyperparameters. The results demonstrate that our method outperforms existing predictors in terms of accuracy for identifying enhancers and their strengths. Importantly, our analyses found that the distribution of bases in enhancers is uneven and the base G contents are more enriched, while the distribution of bases in non-enhancers is relatively even. This result contributes to the improvement of prediction performance and thus facilitates revealing an in-depth understanding of the potential functional mechanisms of enhancers.

**Author summary:** The enhancers contain many subspecies and the accuracy of existing models is difficult to improve due to the small data set. Motivated by the need for accurate and efficient methods to predict enhancer types, we developed a self-attention deep learning model iEnhancer-CLA, the aim is to be able to distinguish effectively and quickly between subspecies of enhancers and whether they are enhancers or not. The model is able to learn sequence features effectively through the combination of multi-scale CNN blocks, BLSTM layers, and self-attention mechanisms, thus improving the accuracy of the model. Encouragingly, by decoupling the CNN layer it was found that the layer was effective in learning the motif of the sequences, which in combination with the self-attention weights could provide interpretability to the model. We further performed sequence analysis in conjunction with the model-generated weights and discovered differences in enhancer and non-enhancer sequence characteristics. This phenomenon can be a guide for the construction of subsequent models for identifying enhancer sequences.

## Introduction

During the development of most eukaryotes, transcription is the first and key step in gene expression of DNA [1, 2]. Enhancers combine transcription factors (TFs), cofactors, and chromatin complexes to act on the promoter to activate or enhance gene transcription [3–6]. In addition, the genetic variation in enhancers was shown to be associated with many human diseases, such as various types of cancer and inflammatory bowel disease [7–14]. Therefore, predicting enhancers becomes a challenging job.

Traditional enhancer prediction is mainly conducted by biological experiments, such as comparing enhancers with TFs [15–17]. However, the biological approach to identify enhancers through experimental techniques is time consuming and inefficient. Thus, many computational methods, such as CSI-ANN [18], ChromeGenSVM [19], rfECS have been proposed in recent years to predict enhancers and mitigation the limitations of experimental methods.

Enhancers are a large set of functional elements consisting of many different subgroups [21], such as strong enhancers, weak enhancers, inhibitory enhancers, and inactive enhancers. Early computational methods are unable to identify the strength of enhancers. For example, EnhancerFinder, GKM-SVM and DEEP [22–24]. iEnhancer-2L [25] is first proposed to solve the enhancer strength prediction problem using a two-layer model. The accuracy is difficult to improve effectively because of the small dataset and insufficient optimization of the base classifier. Despite, many models are proposed to improve the accuracy of predicting enhancers and their strengths such as EnhancerDBN [26], iEnhancer-EL [27], iEnhancer-5Step [28], iEnhancer-ECNN [29] but the accuracy improvement was not significant. The latest model iEnhancer-GAN [30],uses adversarial neural networks to compensate for the significant improvement in enhancer recognition accuracy from small datasets. However, current machine learning methods are still in the black-box learning stage and it is difficult to make interpretable analysis of the model. Although, our previous work iEnhancer-XG [31] have made interpretable analysis of the feature extraction of the model. Nevertheless, there is still much room for improving the interpretability of models and sequences.

In this study, a novel deep learning framework called iEnhancer-CLA is proposed to identify enhancers and their strengths. The model provides a visualization of the model by decoupling the CNN layers to extract position weight matrices (PWMs) and generating the self-attention weights. Provide sequence interpretability by using self-attention weights in combination with other sequence analysis tools, we obtained differences in base combination specificity and base content of different kinds of sequences. The contributions of our model are as follows:

- Construction of a novel multi-scale deep-learning model for identifying enhancers and their strengths. The model could also be extended to predict more enhancer subtypes for future prediction development.
- Using multiscale combination module and Bayesian optimization. Multi-scale modules could extract more sequences of different features and Bayesian optimization could obtain global optimal hyperparameters to improve the efficiency and accuracy of the model.
- Visualization analysis of enhancer sequences and model. The multi-head self-attention layer could generate self-attention weights for each sequence. We find the pattern of bases that are evenly distributed in non-enhancers, in which A and T bases are more enriched, and bases that are not evenly distributed in enhancers, in which base G is enriched that to guide the subsequent drug development.

## Materials and methods

### Benchmark dataset

In the experiments, the dataset is divided into a training set and an independent test set according to the method of Liu et al [25] (Please see the supplementary file, sec.1 for details).

### Model framework

The basic framework diagram of iEnhancer-CLA is shown in Fig 1. The enhancer sequence consists of four nucleotides, namely, adenine (A), guanine (G), cytosine (C), and thymine (T). Enhancer sequences for the four nucleotides (A, C, G, and T) are encoded using one-hot encoding (OHE). The input sequence length is fixed at 200 nt to use the small-batch technique for training and prediction. Therefore, the sequence matrix obtained from the input layer is a fixed-length matrix (4*T, T=200) after using OHE. For the model to learn more sequence features of different lengths, a multi-scale CNN layer with 48 filters of 15 lengths and 48 filters of 25 lengths is applied. With the multi-scale CNN layer being followed by the maximum pooling layer, only the highest value of each four consecutive hidden neurons (pooling length=4 and pooling step=4) is retained in the convolutional layer. The highest value of the layer is retained such that length *T* ^*′*^ = *T*. After the maximum pooling layer filtered out the highest values, the next layer is the BLSTM layer. The sequence length remained the same after BLSTM, and only the encoding size is changed to 32. Lastly, the multi-head self-attention layer is applied. In this frame work, the multi-head self-attention mechanism is used to assess the contribution of sequence regions for localization by multiple heads (head=6), which has the ability to generate self-attention weights for each sequence during the prediction. Stacking of deep learning models at different scales gives the model the ability to adapt to more complex higher order functions. Then, the outputs of the two multi-head self-attention layers are concatenated and fully connected to the output layer, which contains three neurons for the three localization categories. Finally, a softmax activation function is utilized in the output layer, resulting in a prediction value between 0 and 1 for each category.

**Fig 1.**
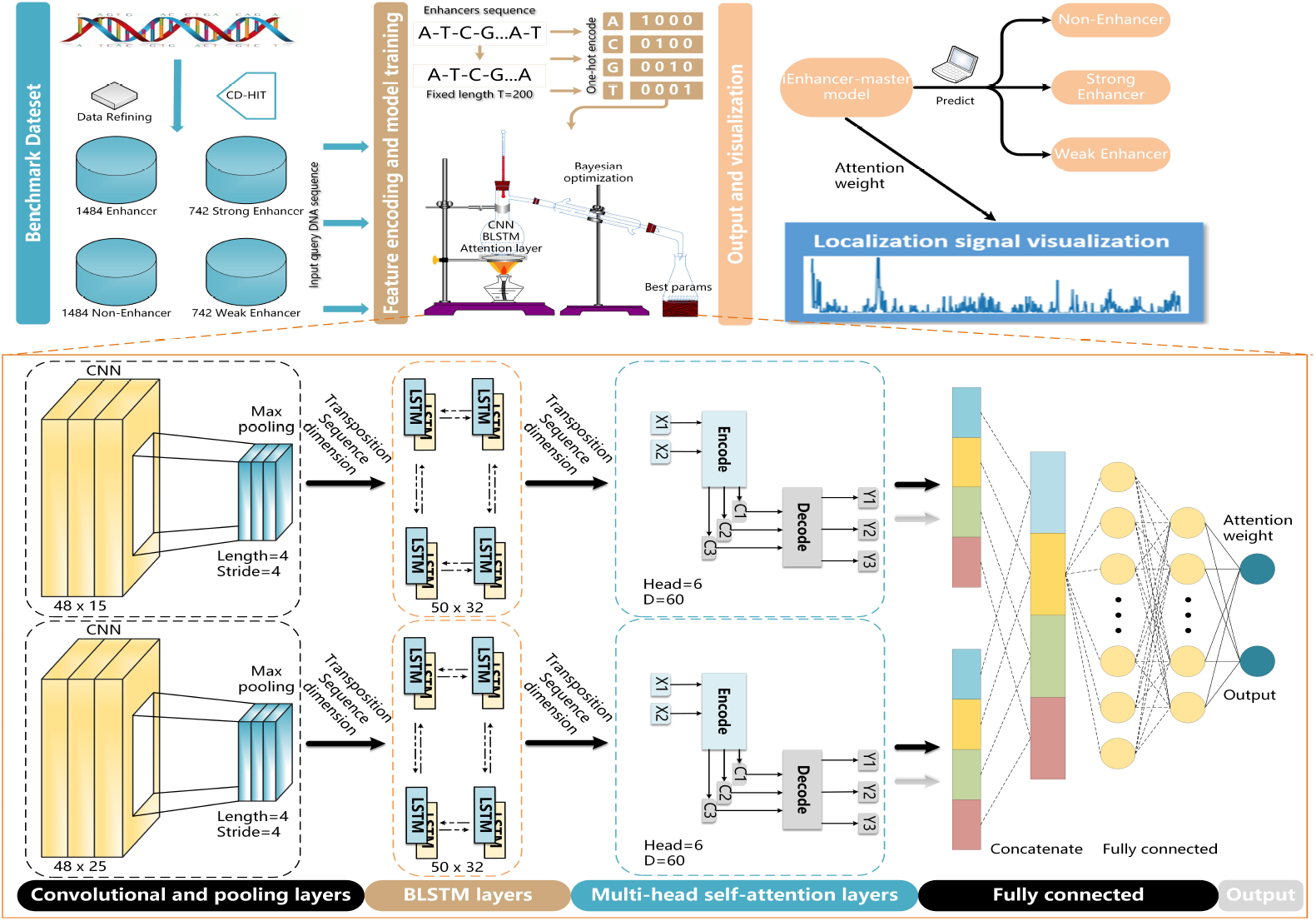
A flowchart of iEnhancer-CLA works. Non-enhancers and enhancers sequences are tagged, sequence fixed length to 200nt and, one-hot encoded. The features are input into the model, and hyperparameters of the model are obtained by Bayesian optimization. Final prediction results of non-enhancers and enhancers can be obtained from the model. The attention weights can be used for visualization of the sequences.

### Multi-head self-attention

The multi-head self-attention model is first established to be used in natural language processing, proposed by [32]. This approach is mainly able to effectively address the problem that the overall semantics of a sentence is composed of multiple components, and the multi-head self-attention model could focus on different parts of the utterance. The method is borrowed in the present work, and the number of *heads* is set to six (derived from hyperparameter tuning). The final weight of the non-enhancer and enhancer are the average weight of the weights obtained for the number of six *heads*.

On the basis of this weight, which part of the enhancer determines whether the sequence is an enhancer or not and the strength of the enhancer could be analyzed. The attention matrix A must first be calculated using Eq (1) to obtain the weights.

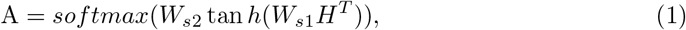

where *H* is the 50-by-32 hidden neuron of the path output obtained from each BLSTM layer; *W*_*s*1_ is a weight matrix of shape *D* − *by* − 32; *D* is the attention dimension hyperparameter, which is 60; *W*_*s*2_ is the matrix of the parameters with shape *heads* − *by* − *D*; and the *head* denotes the number of attention heads, which is 6. *W*_*s*1_ and *W*_*s*2_ are set to the same weight loss values to overcome overfitting and obtain sparse energy scores. The embedding sequence *M*, calculated as a weighted sum by multiplying *A* and *H* as in Eq (2), is also retained for further prediction.

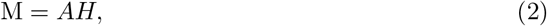

When training the model, the following penalty term *P* in Eq (3) is introduced.

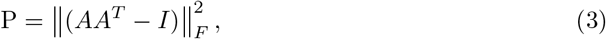

where *A* is the attention matrix, *I* is an identity matrix, and ‖ • ‖ _*F*_ denotes the Frobenius parametrization of the matrix. The greater the similarity between these two attention vectors is, the higher the loss. Thus, by this penalty term, the self-attention model is able to focus on different parts of the enhancer sequence. The loss weight of the penalization term is 0.001.

### Sequence binding motifs visualization in CNN layer

The position weights matrices (PWM) of the sequence binding motifs acquired in each convolutional kernel can be obtained from the CNN filter. Here we follow the approach of Wang et al [33] to obtain the position weights. In the two types of CNN filters (the 15-length, 25-length), we scan each CNN layer to obtain the motifs learned by the convolutional kernel in it. For each convolutional kernel we obtain a set of sequence fragments that have the same length as the convolutional kernel. Finally, we obtained 48 PWMs for 15-length CNN,48 PWMs for 25-length CNN.

### Models and sequences visualization

The model is visualized using a combination of the following two methods. (1) Obtaining PWMs from the CNN layer and comparing them with the motifs of the sequences can prove the validity of the sequence features learned in this layer. (2) Visualize the sequence self-attention weights generated by the self-attention layer to find out the base combinations of the sequence positions corresponding to the highest values of the weights, which can clearly understand the sequence features learned by this layer. Visual images of the distribution of bases at the first and end; gapped motif and ungapped motif of three different sequences, are obtained separately using different methods. The visualization method for the enhancer sequence is as follows:

- The “ggseqlogo [34]” package is applied to visualize the frequency and attention weights of each base at each position, in which the sequence is divided into two segments being the first 50 bases and the last 50 bases of the sequence. This division allows analyzing the attention at the head end and the end.
- Next, the “GLAM2 [35]” method, a tool in MEME suite 5.1.0, to use to find variable-length, gapped motifs to analyze the attention of the sequence in the middle of the enhancer.
- The most recent tool “STREME [36]” was used to discover sequence ungapped motifs. Using this motif as a sequence motif is compared with PWMs obtained in CNN convolution kernels on the “TOMTOM [37]” tool.
- Visualize the sequence self-attention weights generated in the self-attention layer to find the base combinations at the sequence positions corresponding to the highest values in them.

### Parameter tuning and neural network training

The hyperparameters used in the model are the best values obtained by Bayesian optimization, including hidden dimension; number of the attention; dropout rate; etc. In the iEnhancer-CLA model 5 subsets of data sets are obtained by five-fold cross validation respectively and Bayesian optimization is performed on each of the five subsets. The hyperparameters of the sub-model with the highest accuracy are set as the global optimal model. The Adam stochastic optimization method [38] is also used, with a learning rate of 0.001 and a learning rate decay of 5e-5.

The iEnhancer-CLA model uses five-fold cross-validation to dividing the dataset into five parts, four of which are applied for model training and the remaining one for evaluation. The final iEnhancer-CLA model is a collection of five models, and the predictions of the enhancers are the average of the predictions of the five models. For a classification problem, the training goal is to distinguish as little as possible between the prediction vector and the true label vector. The prediction of an enhancers sample is denoted as *x*_*i*_, and its corresponding true label vector is denoted as *y*_*i*_. Each element of *y* is a binary value, denoted as *y*_*ij*_, *j* ∈ [1, 2, 3], indicating whether the enhancer sample belongs to a certain positional category. If *y*_*ij*_ is 1, then *x*_*i*_ belongs to category *j*; otherwise, it belongs to 0. A binary cross-entropy loss function is applied to process each category independently. The loss of sample *x*_*i*_ is defined as follows:

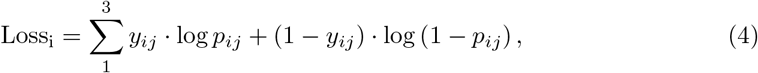

where *p*_*ij*_ ∈ [0, 1]is an element of *p*_*i*_, which represents the prediction vector of sample *x*_*i*_. Given the three locus categories, the loss of sample *x*_*i*_ is the sum of the losses of the individual locus categories. In the construction of this deep learning model, the framework of TensorFlow 1.13.1 and Keras 2.3.0 was used to implement it. The model was trained and tested using GPU Nvidia RTX 2080ti.

## Results and discussion

### Performance evaluation metrics

Here, a five-fold cross-validation method was used to obtain fair data. The area under the ROC curve (AUC), accuracy-recall (PR) curve and Matthew correlation coefficient (MCC) were used to evaluate the performance of the model. Accuracy (ACC), specificity (SP) and MCC were also used to compare the performance with other models (Please see the supplementary file, sec.2 for details).

### Sequence analysis

In this model, we first extracted the first 50 bases at the beginning and 50 bases at the end of the sequence for three kinds of sequences (strong enhancer, weak enhancer, non-enhancer). Using the “ggseqlogo” sequence visualization tool, and the results are shown in Fig 2. The visualization results of the non-enhancer are shown in Figure 2A and 2B. Compared with the enhancer sequence the distribution of bases in the non-enhancer sequence is more uniform, and it can be seen from the figure that the signals of all four bases are obvious. The signal intensity of bases base A and T is higher, which proves that bases A and T are more enriched in the sequences. Strong enhancer sequences are shown in Figure 2C and 2D. It can be seen from the figure that the signal disappears in many positions, which proves that the distribution of bases of different strong enhancer sequences is not specific, and also proves the uneven distribution of bases. From the position of signal, the signal of bases G and C is more significant. Weak enhancer sequences are shown in Figure 2E and 2F, a few positions of the weak enhancer sequences showed the loss of base signals, which proved that the distribution of bases in different weak enhancer sequences was similar. Meanwhile, the signals of bases A and T are more obvious, which proves that bases A and T are more enriched in the sequences.

**Fig 2.**
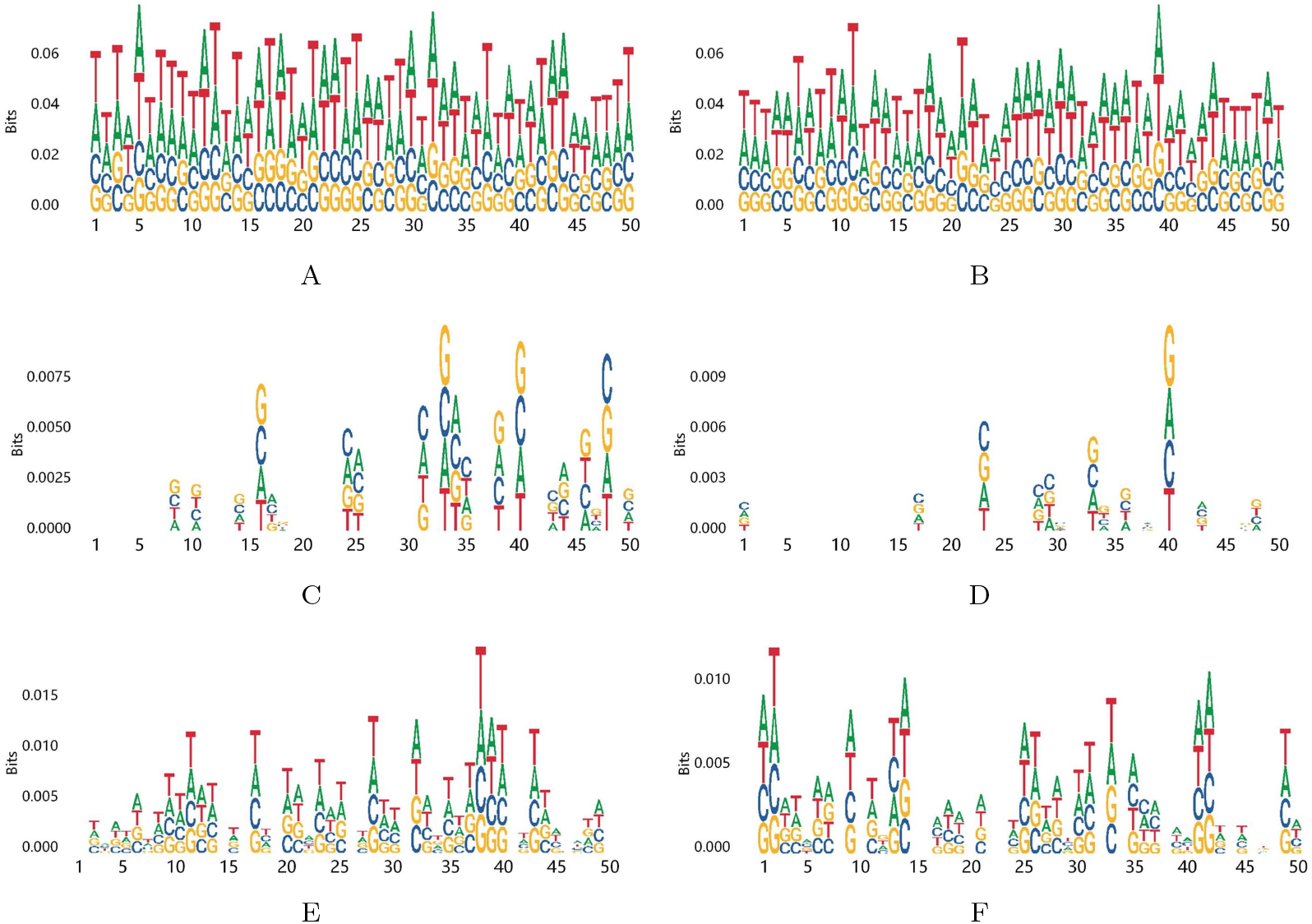
Note the 50-base attention weights near the end of the non-enhancer and enhancer sequences level, which indicates a potential localization signal. (A) First segment of non-enhancer. (B) Last segment of non-enhancer. (C) First segment of strong enhancer. (D) Last segment of strong enhancer. (E) First segment of weak enhancer. (F) Last segment of weak enhancer.

We summarize the visualization results of the three different sequences to obtain the following conclusions: The results from the signal distribution of bases at the first and last ends. The distribution of bases in different sequences in the weak enhancer and non-enhancer sequences were similar. The distribution of bases in different sequences of strong enhancers was in different positions, so signal deletion occurred. From the results of signal intensity of bases. The signal intensities of bases A and T of weak enhancer and non-enhancer were higher, which proved that these two bases appeared more frequently and more abundantly in the sequences. The higher signal intensity of bases G and C in strong enhancers proves that bases G and C are more enriched in strong enhancers.

In “ggseqlog”, the difference in the intensity of the base signals of the enhancers and non-enhancers were found at the first and last ends. Some problems from it were also found, as the base signals of the strong and weak enhancers are not obvious. One possible reason is that the same sequence features do not appear at the same positions. Therefore, the binning signals in the middle of the enhancer sequences were analyzed and visualized using the “GLAM2” tool in the MEME suite, a method that could find the same sequence features at different positions in different sequences. In Fig 3, the localization signals of 300 randomly selected non-enhancer, strong-enhancer, and weak-enhancer sequences were visualized. All parameters were used as default values for the “GLAM2” online tool.

**Fig 3.**
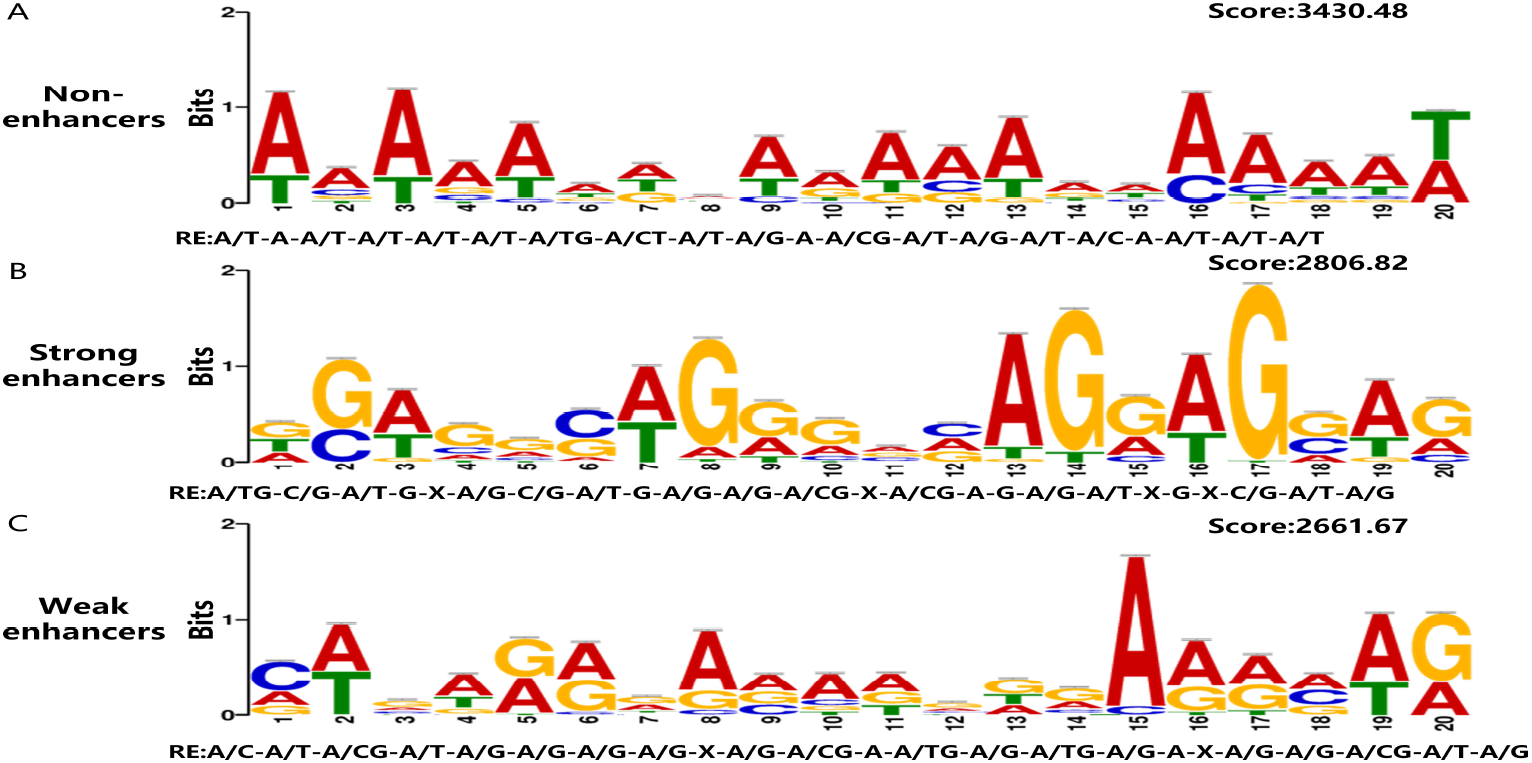
The GLAM2 results for segments from non-enhancers and enhancers sequences. For each result, its rank, score, sequence logo and the regular expression (RE) of the motif are given.

The internal signals of two different enhancers and non-enhancers were obtained in “GLAM2”. As shown in Fig 3, the signal is the same base combination at different positions of different sequences of three kinds of sequences. Regular expression (RE) of the motif refers to the same base combination sequence, where the concept of A/T is that the base at this position could be A or T, while X represents could be any base. The motifs of different sequences in the Fig 3 have their corresponding scores. The higher the score, the more frequent the motif appears in the sequence. We can obtain the base combination characteristics of the three different sequences from Fig 3, which is a guide for the subsequent enhancer prediction.

We have combined the two methods to conclude the following from Fig 2 and 3. The distribution of bases in different sequences in the non-enhancer is similar, with higher intensity of base A and T signals. The most frequent base combination in the non-enhancer was found to be (RE: /T-A/T-A/T-G-A/CT-A/T-A/G-A/CG-A/T-A/T-T) by “GLAM2”. Combining the above findings leads to the conclusion that bases A and T are more enriched in non-enhancers than G and C. The distribution of the bases of different strong enhancer sequences is different, and the signal intensity of bases G and C are higher.

Using “GLAM2”, the base sequence of this sequence was found to be (RE: A/TG-C/G-A/T-G-X-A/G-C/A/G/A/G-A/G-A/CG-X-A/CG-A-G-A/G-A/T-T-A/T-A/G-A). Thus, it is concluded that base G is more enriched in strong enhancers. The different weak enhancer sequences have similar base distribution and higher intensity of base A and T signals. The most frequent base combinations in the weak enhancers were found by “GLAM2” was (RE: A/C-A/T-A/CG-A/G-A/G-A/G-A/G-X-A/CG-A/G-A/G-X-A/G-A/G-A/G-A/G-X-A/G-A/G-A/CG-A/T-A/G). Thereby, the base A is more enriched in the weak enhancer.

### Analysis of CNN motifs

We first used “STREME” tool to analyze the three sequences separately, and obtained 10 ungapped motifs from non-enhancer sequences, 4 ungapped motifs from strong enhancer sequences, and 3 ungapped motifs from weak enhancer sequences. Then, in the multiscale CNN filter, PWMs matrices of 2 lengths (15-length, 25-length) were obtained separately. The PWMs were used to represent the sequence binding motifs. Finally, we use the TOMTOM tool to map the motifs learned from each convolutional kernel to each of the three different sequences motifs obtained in STREME.

The best match in the 15-length CNN filter is obtained as shown in Fig 4 (The top half of the figure is the motif obtained in STREME, and the bottom half is the PWMs extracted from the CNN filter). We can see in Figures 4A and 4B that the CNN extracted PWMs are able to match the two different motifs of the non-enhancer. This also shows that the CNN layer is able to learn useful sequence features in the model. The motifs of the strong and weak enhancers also have matching PWMs respectively. Combining the above 6 sets of comparisons, we can conclude that in this model, the first layer using CNN layer is able to learn many useful sequence features that contribute to the model results.

**Fig 4.**
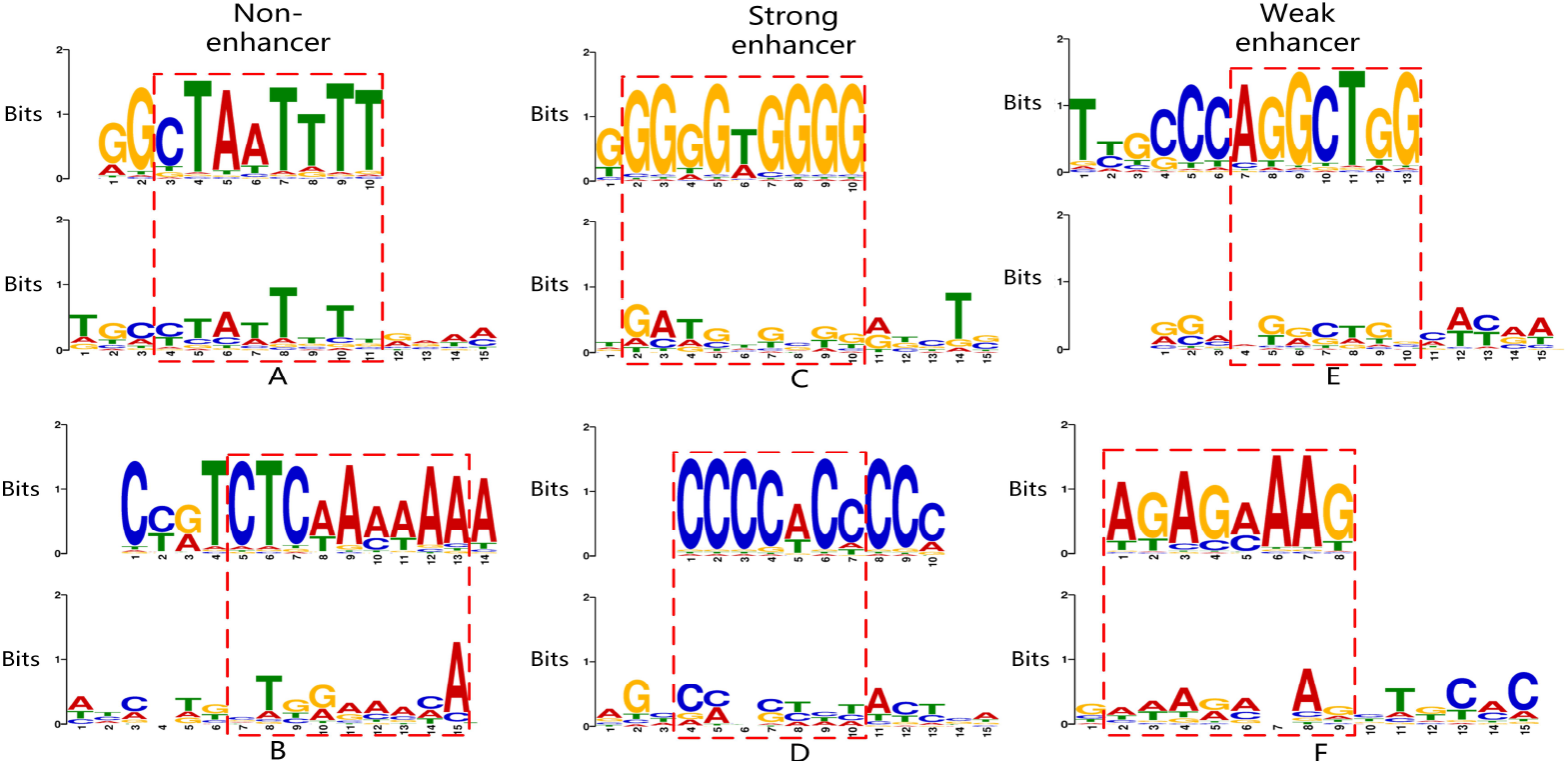
Comparison of the motifs of non-enhancer and enhancer sequences with the PWMs extracted by CNN. (A) and (B) Comparison of different motifs of non-enhancer sequences with different PWMs extracted from CNN. (C) and (D) Comparison of different motifs of strong enhancer sequences with different PWMs extracted from CNN. (E) and (F) Comparison of different motifs of weak enhancer sequences with different PWMs extracted from CNN.

At the same time, the ungapped motif obtained in “STREME” is combined with the results obtained by the previous two tools to further prove our conclusion. The high signal intensity of bases A and T in the non-enhancer motifs in Figures 4A and 4B proves that the content of bases A and T in the non-enhancer sequence is higher than other bases, and the combination of bases A and T is also characteristic of the non-enhancer sequences. Similarly, Figure 4C and Figure 4D show the motif obtained for the strong enhancer in “STREME” shows that the signal intensity of bases G and C is higher than that of other bases. It proves that bases G and C are more enriched in strong enhancers, and the combination of bases G and C is the sequence characteristic of strong enhancers. Motif obtained for the weak enhancer in “STREME” in Figure 4E and Figure 4F shows that the signal intensity of bases A and G is higher than that of other bases. It proves that bases A and G are more enriched in weak enhancers, and the combination of bases A and G is the sequence characteristic of weak enhancers.

### Visualization of attention weights on sequences

In this work, the self-attention layer is able to learn the features of each sequence and generate the self-attention weights of the sequence. With the weights, we can find the differences in base combinations between different kinds of sequences. First, sequences were randomly selected among three different sequences, and the attention weights were visualized separately as shown in Fig 5. Then 10 sequence weights in each sequence are selected separately and averaged as shown in Figure 5D.

**Fig 5.**
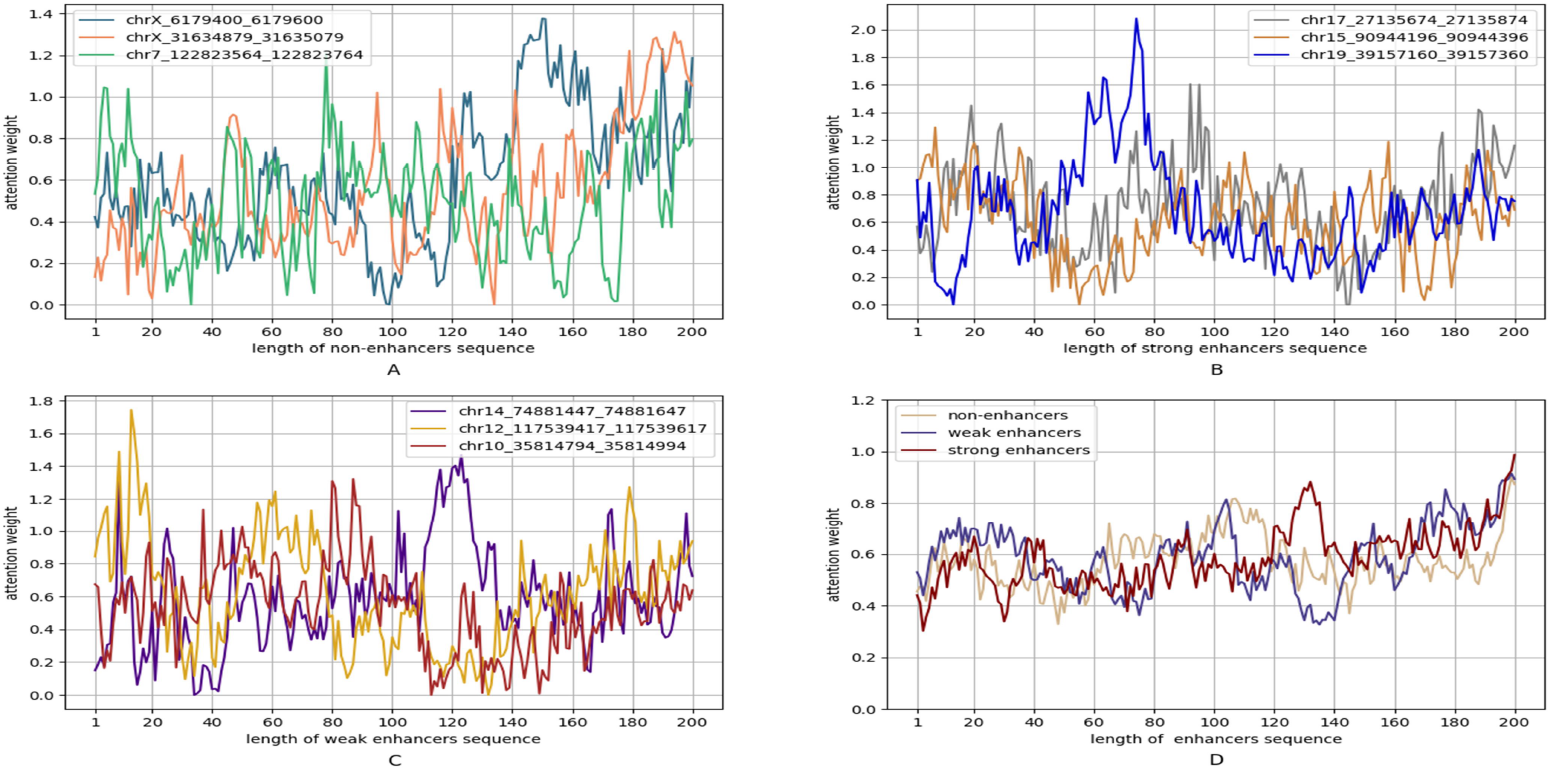
The visualization of attention weights for non-enhancer and enhancers. (A) Random three non-enhancers sequences. (B) Random three strong enhancers sequencers. (C) Random three weak enhancers sequences. (D) Average of 10 sequences of three enhancers.

The base sequences that reached the largest area of the peak region were extracted from the randomly selected sequences to determine what combination of bases could give the sequences higher weights. In Figure 5A, the base sequences of the three non-enhancer highest peaks were extracted as follows:(chrX 6179400)at 140 to 160nt, the base sequence was (TGCAACAACCTGGCTAAGCT); (chrX 31634879) at 180 to 200nt, the base sequence was (AAAAGATGGTACCGGGATGA); and (chr7 122823564) at 70 to 81nt, the base sequence was (TTTGGACTTCT). The results showed that the sequences with consecutive bases of AA or TT or a combination of A and T could reach a peak of attention weight, proving that bases A and T played a key role in non-enhancers. Similarly, the base sequences with the highest peak values of the three enhancer sequences in Figure 5B were extracted. (chr17 27135674) at 90 to 100nt, the base sequence was (CATCACCCAC). (chr15 90944196) at 6 to 12nt, the base sequence was (ACCCCCT). (chr19 39157160) at 66 to 77nt, the base sequence was (CCCGCTCGGAG). The sequences with consecutive C or base combinations of C and G could reach the peak of attention weight, proving that C and G played a key role in the strong enhancer. The highest peak base sequences of the three sequences in the weak enhancer are as follows: (chr14 74881447) at 118 to 127 nt, with the base sequence (GTGGTGTGTGT); (chr12 117539417) at 10 to 20 nt, with the base sequence (GGGGATAGG); and (chr10 35814794) at 80 to 90 nt, with the base sequence (TGCATTCTAT). From the base combinations of the three sequences at the highest peak, consecutive G combinations of base sequences that could reach the higher peak were found. The combination of A and T was still present, but G appeared more frequently among them, thus proving that G plays a key role in the weak enhancer. The above results demonstrated that the combination of the content of A and T bases played a dominant role in non-enhancers, while the content and combination of G bases played a dominant role in enhancers.

The average values of the weights for the three kinds of sequences are shown in Figure 5D. Although the general trends are the same, some differences exist between each kind of sequence. When the sequence position is at 1 to 30nt, the weight of weak enhancer is significantly higher than the others. At 60 to 80nt, the weight of non-enhancers is higher than that of enhancer sequences. At 120 to 140nt, the weight of the strong enhancer is higher than the other kinds of sequences, and the weight has an obvious upward trend, while the weak enhancer has an obvious downward trend in this segment. This result shows that the main differences between strong enhancer, weak enhancer, and non-enhancer sequence features are concentrated in three positions (1-30nt,60-80nt,120-140nt). This finding also serves as a guide for our subsequent analysis and prediction of enhancer sequences.

The PWMs extracted in the CNN layer and the sequence self-attention weights generated in the self-attention layer to see the model can learn the sequence features of different sequences to contribute to the prediction. We can also clearly understand what sequence features are learned in the first layer of the model and what sequence features are learned in the last layer of the model, thus achieving interpretability of the model.

Also, the base sequence corresponding to the highest value of self-attention weight for each sequence combined with the results of the previous three methods were able to demonstrate the differences in base combination and base content of the three different sequences.

### Comparison between iEnhancer-CLA and existing methods and tools

Given that the proposed model is a first single-layer multiclassification enhancer prediction model, comparing it with other existing two-layer models was difficult. Thus, several classical enhancer two-layer models were selected for comparison.

The proposed method was compared with the classical method on the same independent-test dataset to objectively evaluate the prediction performance. The following four evaluation metrics were also selected for evaluation: ACC, AUC, SP, and MCC.

As shown from the results in Table 1, although the work was multi-classified, the prediction results were still higher than those of classical model methods under the chosen evaluation metrics. In particular, for the prediction of strong and weak enhancers, the ACC values were significantly higher than those of the previous classical models. The established model was able to perform multi-classification prediction in a single layer, and it showed better accuracy than the previous classical models.

**Table 1.** Comparison of the proposed predictor with the state-of-the-art predictors in identifying enhancers (the first layer) and their strength (the second layer) on the independent dataset.

|  | Method | ACC% | AUC% | SP% | MCC | Source |
| --- | --- | --- | --- | --- | --- | --- |
| First layer | EnhancerPred | 74.00 | 80.10 | 74.50 | 0.480 | Jia and He, 2016 |
|  | iEnhancer-2L | 73.00 | 80.60 | 75.00 | 0.460 | Liu et al., 2016 |
|  | iEnhancer-EL | 74.75 | 81.70 | 78.50 | 0.496 | Liu et al., 2018 |
|  | iEnhancer-ECNN | 76.90 | 83.20 | 75.20 | 0.537 | Nguyen et a., 2019 |
| iEnhancer-CLA | Non-enhancer | **77.00** | **83.62** | **79.00** | **0.540** | This study |
| Second layer | EnhancerPred | 55.00 | 57.90 | 65.00 | 0.102 | Jia and He, 2016 |
|  | iEnhancer-2L | 60.50 | 66.80 | 74.00 | 0.218 | Liu et al., 2016 |
|  | iEnhancer-EL | 61.00 | 68.00 | 68.00 | 0.222 | Liu et al., 2018 |
|  | iEnhancer-ECNN | 67.80 | 74.80 | 68.00 | **0.368** | Nguyen et a., 2019 |
| iEnhancer-CLA | Strong enhancer | **78.50** | **84.69** | **91.33** | 0.365 | This study |
|  | Weak enhancer | **74.25** | 63.16 | **96.66** | 0.078 | This study |

### Effectiveness of iEnhancer-CLA

Due to the previous two-layer model, one layer was used to distinguish whether it is an enhancer, and one layer was used to distinguish the strength of the enhancers. This computational method, after years of research and development, was able to distinguish enhancers more accurately than before. However, the two-layer model also limits the identification of enhancers. Many subtypes of enhancers exist, not only strong and weak enhancers. In future research, the role of more subtypes of enhancers and the use of computational methods to distinguish and predict must be developed and studied. The importance of the proposed multi-labeled single-layer multiclassification prediction model was remarkably demonstrated in the present work, of which the results in Table 2 were obtained using five-fold cross-validation. Table 2 shows the results for the training set. We put the results of the independent test set and the results of the non-enhancer sequences and enhancer sequences balanced data set in the (supplementary file, sec.3).

**Table 2.** The performances of model on train dataset.

|  | ACC% | AUC% | SP% | MCC |
| --- | --- | --- | --- | --- |
| Non-enhancer | 79.85 | 88.57 | 0.8793 | 0.5972 |
| Strong enhancer | 82.21 | 87.71 | 0.6793 | 0.4902 |
| Weak enhancer | 76.11 | 72.64 | 0.4712 | 0.2019 |

## Discussion and conclusion

With the rapid development of computer methods, this work focused on how to use efficient methods to identify enhancer sequences and enhancer the strength. Previous works had many kinds of methods to identify enhancers. For the prediction model of single-layer multiclassification with multiple labels proposed here, the same previous dataset was used, but each kind of sample in the dataset with labels was annotated. In the experiments, a five-fold cross-validation dataset was built from the benchmark dataset. A BLSTM layer and a novel multi-head self-attention mechanism were added in the proposed model on the basis of the CNN model. The proposed model could identify enhancers more accurately than others, as indicated by the results of the independent-test set.

The model was able to generate sequence-related attention weights and visualize the sequences by using sequence analysis software with “ggseqlogo”, “GLAM2” and “STREME”. The differences between the two kinds of enhancers and non-enhancers were identified, which could be useful for future research on enhancers. The combination of the PWMs extracted in the CNN layer and the self-attention weights in the self-attention layer provides interpretability for the model. The results changed the previous black box experiment and improved the credibility of the experiment.

